# ISL2 is an epigenetically silenced tumor suppressor and regulator of metabolism in pancreatic cancer

**DOI:** 10.1101/2020.05.23.112839

**Authors:** Turan Tufan, Jiekun Yang, Krishna Seshu Tummala, Harun Cingoz, Cem Kuscu, Sara J. Adair, Gamze Comertpay, Sarbajeet Nagdas, Bernadette J. Goudreau, Husnu Umit Luleyap, Ku-lung Hsu, Dave F. Kashatus, Nabeel Bardeesy, Todd W. Bauer, Mazhar Adli

## Abstract

Pancreatic ductal adenocarcinoma (PDAC) remains one of the deadliest cancers. Uncovering mechanisms responsible for the heterogeneous clinical features of this disease is an essential step toward developing improved and more specific therapeutic approaches. Here, we sought to identify transcriptional regulators of aggressive PDAC growth through *in vivo* CRISPR screening of epigenetic and transcription factors in an orthotopic model. We identified the ISL LIM homeobox 2 (**ISL2**) gene as a tumor suppressor whose depletion enhances the proliferation of human PDAC cells *in vitro* and *in vivo* and cooperates with activated KRAS to initiate PDAC in a murine model. Conversely, the upregulation of *ISL2* expression through CRISPR-mediated locus-specific epigenetic editing results in reduced cell proliferation. Importantly, *ISL2* is epigenetically silenced through DNA methylation in ~60% of PDAC tumors, which correlates with poor patient outcome. Functional studies showed that ISL2 loss rewires metabolic gene expression, and consequently potentiates oxidative phosphorylation while reducing glycolysis. This metabolic shift creates selective vulnerability to small molecule inhibitors of mitochondrial respiration and fatty acid oxidation. Collectively, these findings reveal ISL2 as a novel tumor suppressor whose inactivation drives metabolic reprogramming in an aggressive PDAC subset and point to potential therapeutic vulnerabilities in these tumors.

## INTRODUCTION

Pancreatic ductal adenocarcinoma (PDAC) is projected to be the second leading cause of cancer death in the USA by 2030^1^. The median survival rate for advanced PDAC is ~6 months, and less than 8% of patients survive beyond five years^2^. Recurrent genomic alterations in PDAC include oncogenic *KRAS* mutations (>90% of tumors)^3^ and loss of function mutations in the *CDKN2A*, *TP53*, and *SMAD4* tumor suppressors (~40-70%). Unfortunately, none of these common genetic alterations is currently targetable, and attempts to inhibit KRAS effectors have been largely unsuccessful^4–6^. Moreover, the mutational profiles of these genes do not fully explain the heterogeneous transcriptional signatures and clinical features observed across different patients with the disease.

The transcription factors and chromatin regulators that control PDAC phenotypes in vivo remain incompletely defined. Recent studies have documented transcriptional programs governing metastasis and the identity of different histological subtypes of PDAC^7–13^. Here, we sought to identify new transcriptional pathways dictating aggressive PDAC growth by performing an unbiased CRISPR-mediated gene knockout (KO) screening in a clinically-relevant orthotopic patient-derived xenograft (PDX) model. We discovered *ISL2*, a relatively uncharacterized transcription factor, as a top-scoring gene whose depletion enhances *in vitro* cell proliferation and *in vivo* tumor formation capacity of PDAC cells. We also found that ISL2 loss drives a distinct metabolic program in this setting and demonstrated that epigenetic silencing of ISL2 is frequent in human PDAC patient samples, associated with a concordant metabolic gene expression profile and poor prognosis. Our findings reveal ISL2 loss as a marker and driver of aggressive PDAC growth and suggest new therapeutic opportunities targeting metabolism in this defined disease subset.

## RESULTS

To identify transcription factors and chromatin regulators that restrain PDAC growth in vivo and are potential tumor suppressors, we performed CRISPR gene KO screening in a clinically relevant PDX model in which a patient’s tumor is propagated orthotopically within the pancreas of athymic nude mice^14^. The initial screening was performed in the PDX366 model (mutant for *KRAS*, *P53*, and *SMAD4*)^15^. Short-term cultures of PDX366 cells were engineered to express CRISPR/Cas9 and then infected with viruses containing a “nuclear” sgRNA library targeting ~4,000 human epigenetic regulators, transcription factors, and nuclear proteins^16^. By using a low (~0.25) multiplicity of infection (MOI), each cell was infected with a single sgRNA. After a week of puromycin selection, ~8 million cells (~200× sgRNAs coverage) were injected into the pancreas of mice (n=3; **Figure 1A**). We sought to take advantage of the pooled screen format to identify novel tumor suppressors via identifying sgRNAs enriched in the final tumor population *in vivo*. Accordingly we allowed tumors to form for five weeks and assessed the abundance of each sgRNA in the tumors relative to the day 0 control samples (see **Methods**).

**Figure 1:**
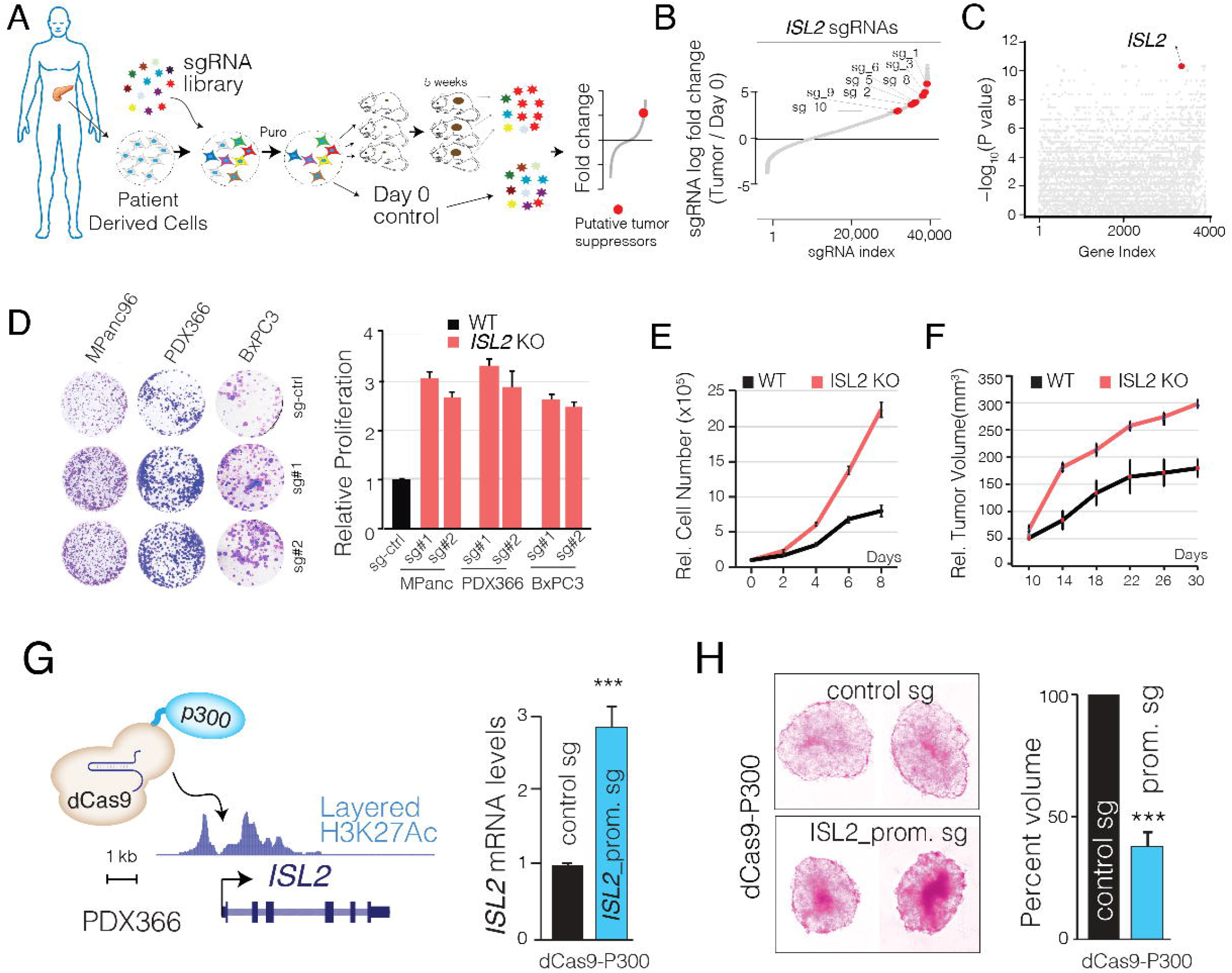
in vivo CRISPR screening identifies ISL2 as a PDAC tumor suppressor. **A)** Schematic showing in vivo CRISPR screening approach**. B)** Dot plot showing enrichment and depletion of all sgRNAs in tumors compared to Day 0 samples. ISL2 targeting sgRNAs are red labeled. **C)** The dot plot shows the gene-level statistical significance of enriched genes. **D)** Crystal violet and bar plots show the relative proliferation of control and sgRNA infected cells after two weeks. **E)** The line plot shows MTT-measured cell proliferation after 8 days of continuous culturing. **F)** The line plot shows *in vivo* tumor volumes for control and ISL2 KO cells in a xenograft model. **G)** Schematic showing recruitment of dCas9-P300 complex to activate endogenous *ISL2* locus. The bar plot shows relative ISL2 mRNA levels in control sgRNA and ISL2-promoter targeting sgRNA infected cells. **H)** Images show 3D-spheres for control sgRNA and ISL2-promoter targeting sgRNAs in cells containing dCas9-p300 complex. The bar plot shows relative sphere volumes in control and ISL2-upregulated cells. Error bars show SEM of three or more independent experiments.

### Loss of ISL2 results in increased PDAC cell proliferation in vitro and enhanced tumor growth in vivo

The most significantly enriched sgRNAs targeted the *ISL2*, *CREM*, and MBD1 genes. These genes scored as top hits based on multiple metrics: individual top-scoring sgRNAs, overall significance, and gene viability score (**Figure 1B, C, Supplementary Figure 1A, B**). Among these hits, ISL2 stood out with 9/10 sgRNAs targeting *ISL2* showing enrichment of >16-fold across the three tumors (Kolmogorov-Smirnov test p-value < 1.10^−10^; **Figure 1B-C**). *MBD1* has previously been identified as a negative regulator of cancer cell proliferation^17^, whereas neither *ISL2* or *CREM* are characterized in this context. We conducted initial validation studies by generating two independent knockout lines for either ISL2 or CREM and examining proliferation in vitro. While knockout of either gene resulted in significantly higher cell proliferation rates, the effect was most dramatic with *ISL2* inactivation (**Supplementary Figure 1C**)

Encouraged by these results, we focused on *ISL2* and included additional PDAC cell lines in our analyses (MPANC-96 and BxPC3, in addition to PDX366). Independent *ISL2*-targeting sgRNAs diminished ISL2 protein levels more than 90% in a mixed cell population (**Supplementary Figure 1D**), resulting in significantly higher cell proliferation as measured by crystal violet colony formation (**Figure 1D**) and 8-day-MTT cell viability assays (**Figure 1E**). Moreover, we found that ISL2 KO cells grew more efficiently as 3D spheroids (**Supplementary Figure 1E**). Importantly, these effects were recapitulated in vivo, with ISL2 KO MPANC96 cells forming more rapidly growing subcutaneous tumors compared to ISL2 WT MPANC96 cells (**Figure 1F**).

### The upregulation of endogenous ISL2 impairs cellular proliferation

The above studies validated our CRISPR screening data and showed that ISL2 loss provides PDAC cells a growth advantage *in vitro* and *in vivo*. We next assessed whether the upregulation of *ISL2* has the opposite effect. Indeed, the transient transfection of *ISL2* resulted in significant apoptotic cell death. We, therefore, extended these findings by using CRISPR-based epigenetic editing to up-regulate the expression of *ISL2* from the endogenous locus. To this end, we took advantage of the dCas9-P300 approach, where the histone acetyltransferase P300 is recruited to the target loci to locally deposit an H3K27ac activating mark and induce the expression of the target gene^18,19^. Recruitment of dCas9-P300 to the *ISL2* promoter increased *ISL2* mRNA by ~3-fold (**Figure 1G**). This low-level induction significantly reduced cell proliferation as assessed by the 3D sphere assay (**Figure 1H**). We also noted a necrosis-like phenotype in the center of spheres and an increase in apoptosis (**Figure 1H**, **Supplementary Figure 1F**). Thus, PDAC cells are highly sensitive to modulation of ISL2 levels, and even modest increases in ISL2 expression markedly attenuates the growth of PDAC cells.

### Expression of ISL2 reduces during PDAC development through DNA methylation

Our functional data predict that there may be selective pressure to inactivate ISL2 in human PDAC. We, therefore, analyzed existing public data sets for evidence of genetic or epigenetic alterations of ISL2 in PDAC. While we did not observe recurrent mutations or deletions, we found *that ISL2* mRNA expression is significantly down-regulated in primary PDAC tissue compared to matched normal tissue in both ICGC^20^ and TCGA projects^21^ (**Figure 2A).** PDAC originates from precursor lesions, most commonly pancreatic intraepithelial neoplasms (PanINs) and intraductal papillary-mucinous neoplasms (IPMNs). IPMN arises from mucinous adenomas (IPMA), which progress into mucinous carcinoma (IPMC) and eventually into PDAC. We also used existing data sets from laser-captured microdissected pancreatic neoplasms (IPMA, IPMC, and PDAC) at different stages (GSE19650)^22^ to test whether ISL2 expression is down-regulated during PDAC progression. The analysis indicates that ISL2 expression is significantly down-regulated in the IPMN stage, suggesting that loss of ISL2 expression is likely an early event during PDAC development (**Figure 2A**).

**Figure 2:**
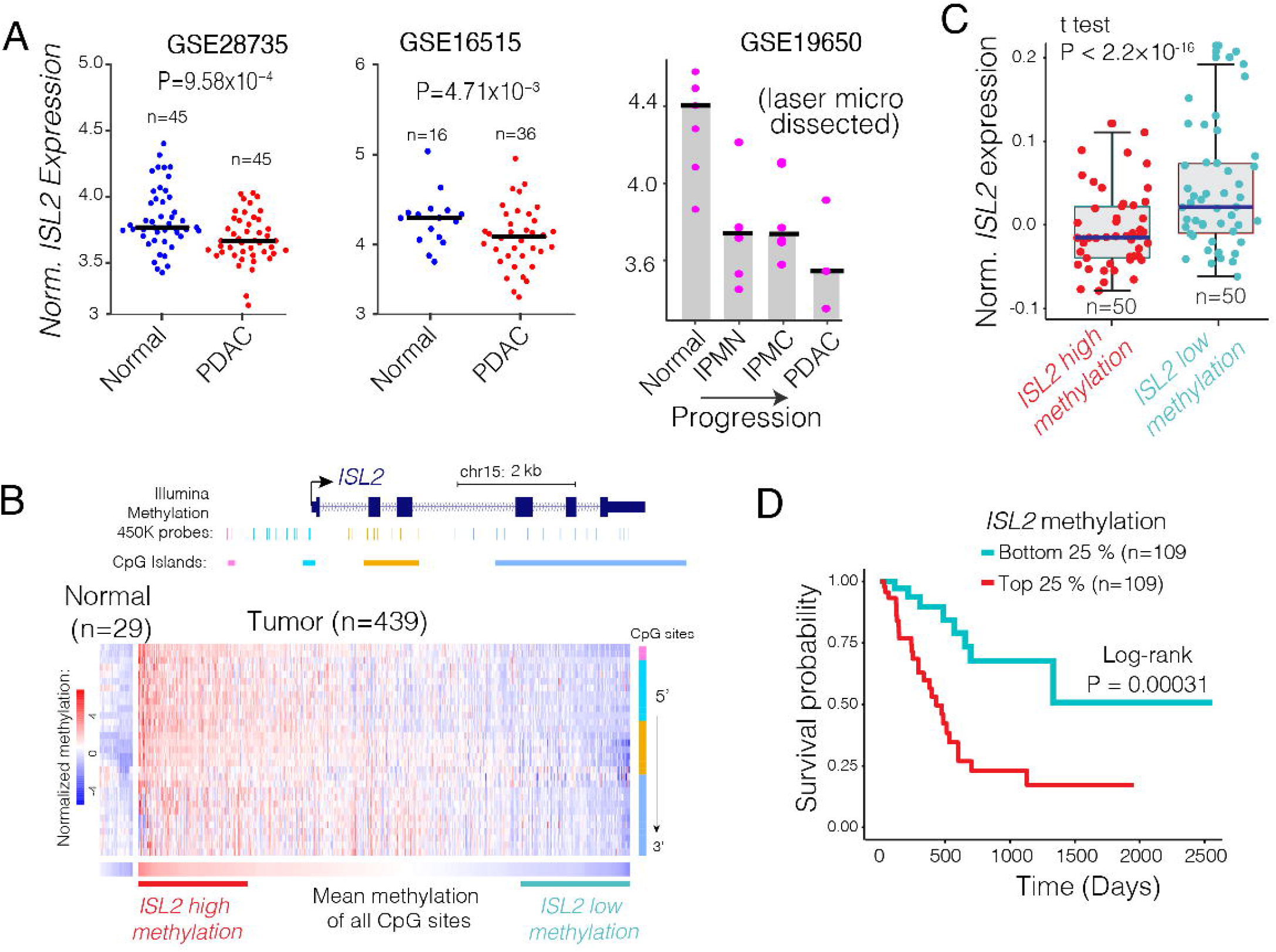
ISL2 is epigenetically silenced and downregulated in primary PDAC tumors. **A)** Dot plot shows ISL2 mRNA levels in multiple independently generated normal and matched primary PDAC tumors. **B)** Heatmap shows relative DNA methylation levels across all CpG islands at ISL2 locus in the normal pancreas (n=29) and primary PDAC tumors (n=439). C) Box plots show the relative ISL2 mRNA levels in the top and bottom 25% methylated primary PDAC tumors. D) Kaplan-Meier survival plots showing overall survival probability for patients having methylated (red, top 25%) and unmethylated (blue, the bottom 25%) ISL2 tumors.

Importantly, we found that the *ISL2* locus is hypermethylated in a significant fraction of PDAC tumors (**Figure 2B**). Specifically, when we compared the methylation levels at all 31 CpG sites at ISL2 locus, we observed that more than 60% of primary PDAC tumors had significantly higher methylation levels compared to normal tissues (student’s t-test p < 1×10^−13^) (**Figure 2B**). DNA methylation is often associated with gene repression^23^. Correspondingly *ISL2* hypermethylation was strongly correlated with reduced levels of *ISL2* expression (**Figure 2C**). Moreover, *ISL2* hypermethylation in resected specimens was associated with reduced overall survival (log-rank test p = 0.00031; **Figure 2D**). These findings support the significance of our screening results and indicate that *ISL2* silencing by DNA methylation during PDAC progression results in more aggressive tumors.

### ISL2 inactivation in pre-malignant PanIN organoids accelerates PDAC development and invasiveness in vivo

Mutational activation of KRAS is an early event in PDAC pathogenesis, serving to trigger the formation of premalignant PanIN lesions. Notably, the integrative gene expression analysis of more than 300 PDAC tumors highlights that KRAS expression levels is inversely correlated with ISL2 expression **(Figure 3A).** Hypothesizing that loss of ISL2 is a critical molecular event downstream of oncogenic KRAS signaling, we next examined the potential cooperative interactions between ISL2 loss and KRAS^G12D^ in PDAC progression using an organoid model derived from murine PanIN-derived organoids (from Pdx1^Cre^:Kras^LSLG12D^) ^24,25^. We generated stable Kras^LSLG12D^ organoid lines with two different lentiviruses expressing sgRNAs that target ISL2 or with a control guide targeting bacterial LacZ. Following verification of CRISPR-mediated ISL2 ablation, we injected each organoid line subcutaneously into the flanks of SCID mice **(Figure 3B).** Examination at 2-month post-injection revealed that the ISL2 depleted organoids formed significantly larger tumors compared to the controls **(Figure 3C).** Histological analysis demonstrated that all the tumors had PDAC histology, although the sgISL2 tumors were notably more invasive (peritoneal invasion was observed in 3/8 sgISL2 tumor and 0/4 sgLacZ tumors; **Figure 3D, E)**. These results reveal ISL2 as a conserved suppressor of PDAC growth.

**Figure 3:**
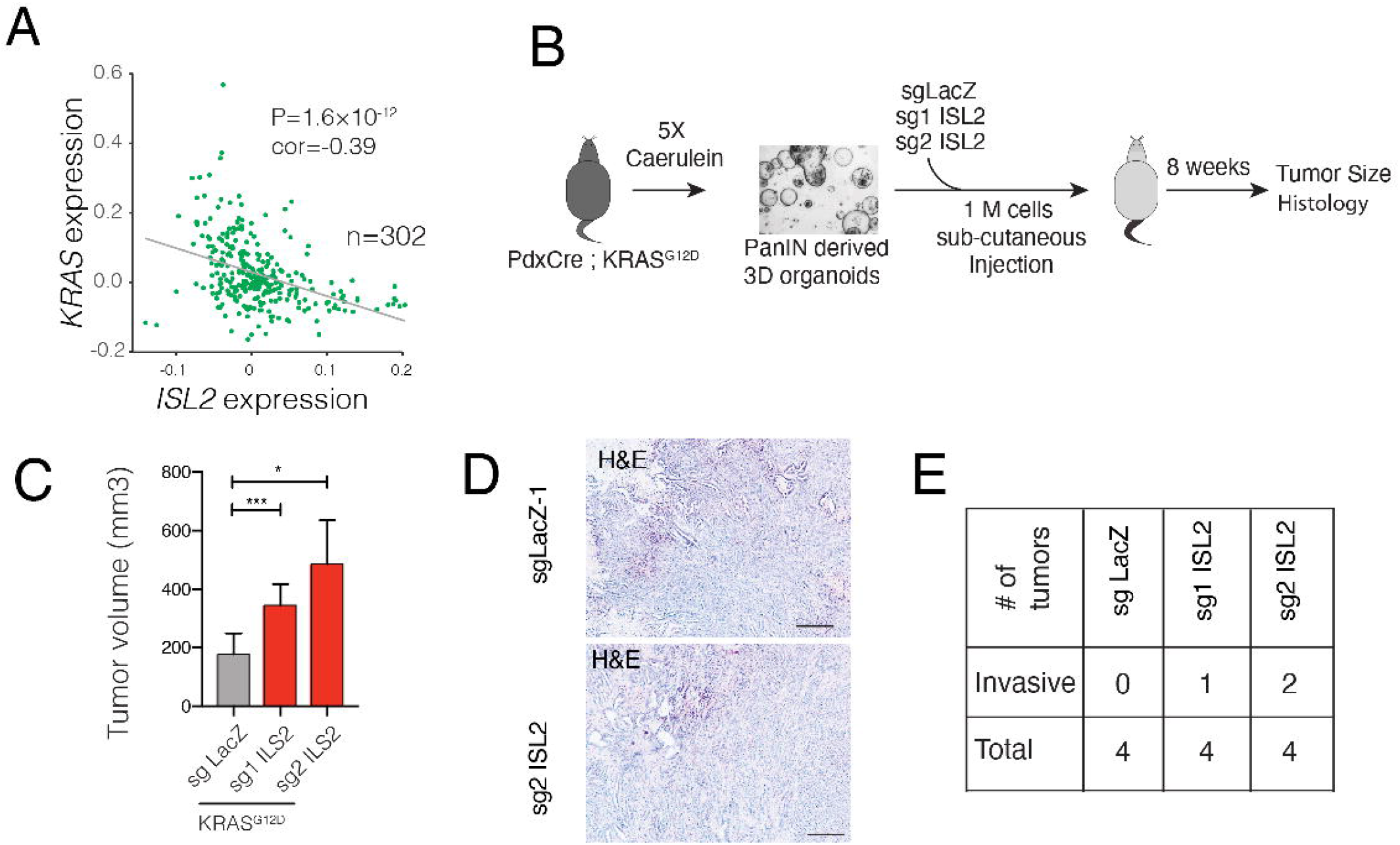
ISL2 silencing cooperates with mutant Kras to drive PDAC progression. **A)** The dot plot shows the relative expression of *KRAS* and *ISL2* genes across a panel of PDAC tumors. **B)** Schematic showing use of murine PanIN-derived organoid model to examine the impact of ISL2 inactivation on tumorigenicity. Organoids derived from Pdx1-Cre;LSL-KRAS^G12D^ mice bearing PanIN lesions were subject to CRISPR-CAS9-mediated gene editing using sgRNAs against ISL2 or LacZ. Organoids (4/group) were tested for tumorigenicity in C57Bl/6 mice. **C)** Bar plots showing relative tumor volumes of organoids expressing the indicated sgRNAs. **D)** Representative hematoxylin and eosin (H&E) staining showing pancreatic ductal adenocarcinoma histology of the tumors formed by the organoids. **E)** Chart showing the frequency of invasion into the peritoneum.

### ISL2 depletion results in expression alterations of metabolic genes

We next set out to understand the molecular changes associated with the depletion of ISL2 and to define the potential mechanisms resulting in the increased proliferation of PDAC cells. Comparison of gene expression in WT and ISL2 KO MPANC and PDX366 cells via RNA sequencing analysis revealed a comparable number of upregulated and downregulated genes (2,303 and 2,224, respectively) upon ISL2 depletion in the two independent PDAC cell lines. Gene ontology (GO) enrichment analysis of differentially expressed genes pointed to potential molecular mechanisms underlying ISL2-mediated cellular phenotypes. Specifically, ISL2 loss was associated with enrichment of OXPHOS and lipid metabolism gene signatures and downregulation of cytoskeletal, mTOR signaling, and glycolysis signatures (**Figure 4A**). The qRT-PCR analysis confirmed that ISL2 depletion reduced the expression of genes with established roles in glycolysis (n = 13) and upregulated OXPHOS genes (n = 10) (**Figure 4B**).

**Figure 4:**
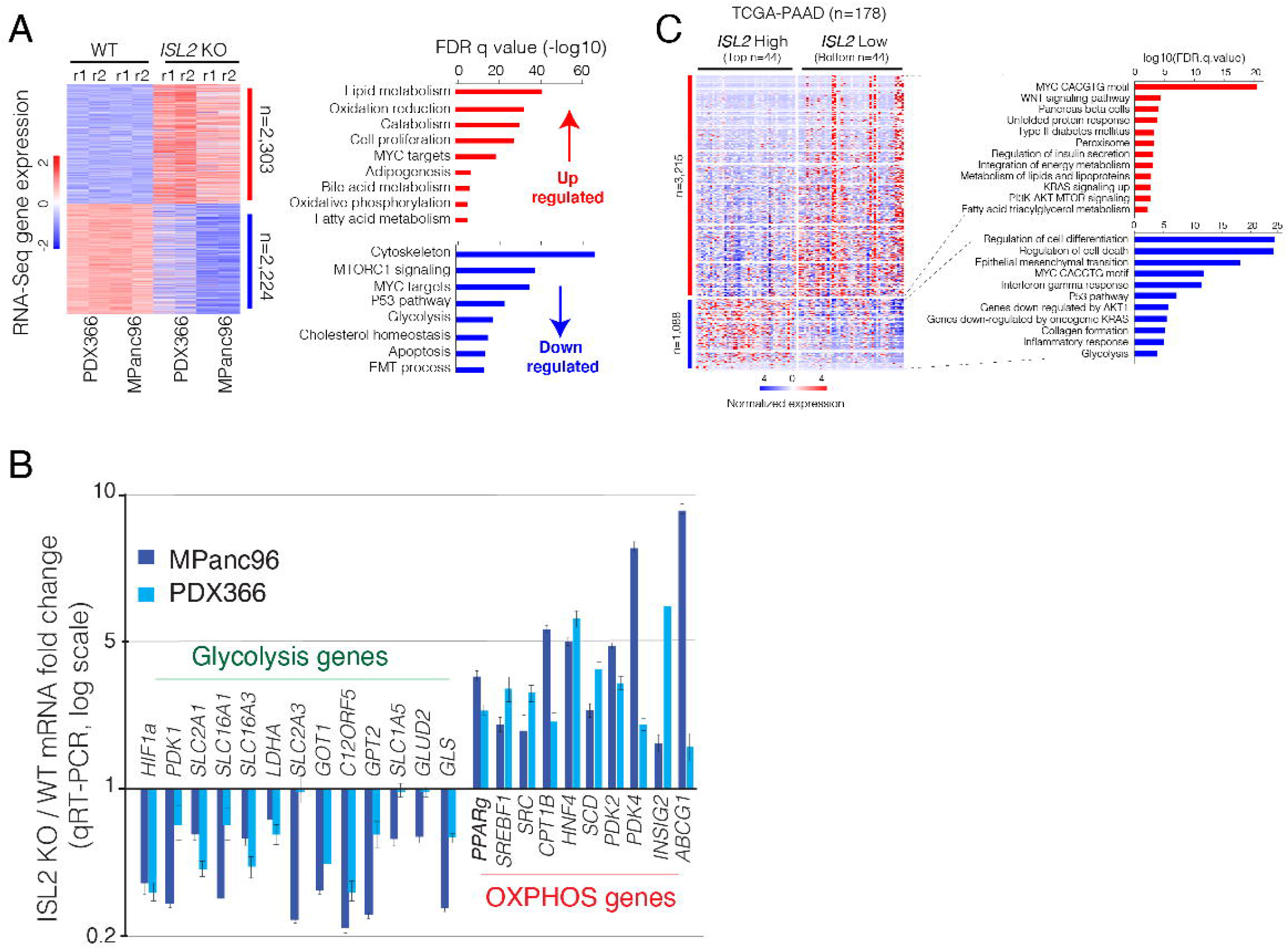
ISL2 regulates the expression of metabolic genes in PDAC cells. **A)** Heatmap shows differentially expressed genes in two replicates of WT and ISL2 KO PDX366 and MPanc96 cells. Bar graphs show gene ontology enrichment levels of genes upregulated (red) and downregulated (blue) in ISL2 KO cells. **B)** Heatmap shows differentially regulated genes in primary PDAC tumors with high (top n=44) and low (bottom n=44) ISL2 expression. Bar graphs show gene ontology enrichment levels of genes in upregulated (red) and downregulated in ISL2 KO cells. **C)** Bar plot shows the qRT-PCR measured expression of genes implicated in glycolysis (n=13) and oxidative phosphorylation/TCA cycle. Error bars show SEM of three or more independent experiments.

We then investigated whether comparable signatures are observed in primary PDAC tumors with differential *ISL2* expression. To this end, we used TCGA RNA-Seq data to stratify the 178 PDAC tumors into high-*ISL2* vs. low-*ISL2* expressing groups (top 44 vs. bottom 44). Consistent with the in vitro data, we found that the primary tumors expressing low *ISL2* have higher expression of genes involved in lipid and fatty acid metabolism and down-regulation of glycolysis-related genes **(Figure 4B).** Thus, the data support the physiological relevance of our in vitro results and indicate that ISL2 status serves as a switch between two distinct metabolic gene expression programs in PDAC.

### ISL2 KO cells have reduced glycolysis but increased OXPHOS

These findings led us to predict that ISL2 depletion results in metabolic reprogramming, shifting PDAC cells to become more reliant on OXPHOS than glycolysis (**Figure 5A**). To test this, we performed functional studies to detect potential metabolic reprogramming due to ISL2 loss. Interestingly, despite showing more rapid growth rates compared to ISL2 WT cells, ISL2 KO cells showed a delay in acidifying the tissue culture media as reflected by the media color. In line with this, a 10-day old culture media from the KO cells contained significantly lower levels of lactate **(Figure 5B)** and higher levels of glucose **(Figure 5C)**, suggesting a reduced glycolytic rate.

**Figure 5:**
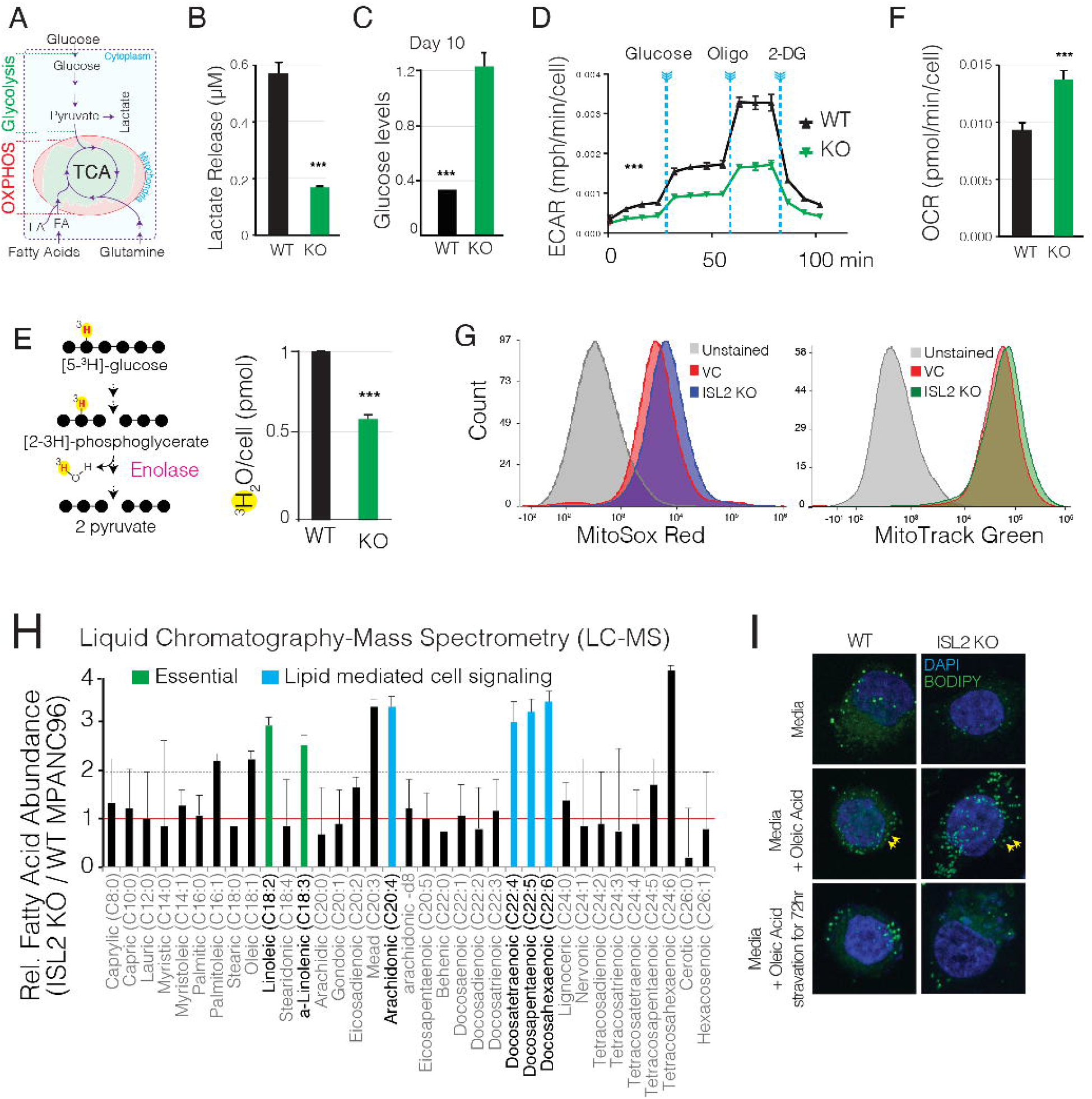
ISL2 depletion reprograms PDAC metabolism. **A)** Schematic showing overall Glycolytic and oxidative phosphorylation (OXPHOS) metabolic reactions. **B)** Bar plots show lactate levels in WT and ISL2 KO MPanc96 cells. **C)** Bar plots show glucose levels in WT and ISL2 KO MPanc96 cells. **D)** The line plot shows Seahorse-measured extracellular acidification rates (ECAR) in WT and ISL2 KO MPanc96 cells. **E)** Schematics show the release of tritium water during glycolysis. Bar plots show relative levels of tritium water in WT and ISL2 KO MPanc96 cells. **F)** Bar plot shows Seahorse-measured oxygen consumption rates in WT and ISL2 KO MPanc96 cells. **G)** Flow cytometry measured Mitosox, and Mitotrack staining levels show relative mitochondrial content and activity, respectively, in WT and ISL2 KO cells. **H)** Bar plot shows liquid-chromatography-mass spectrometry measured relative fatty acid levels in WT and ISL2 KO cells. **I)** Fluorescent-microscopy images show BODIPY measured lipid droplet content in WT and ISL2 KO cells. Error bars show SEM of three or more independent experiments.

We used a series of metabolic assays to more directly examine the impact of ISL2 loss on PDAC cell metabolism. First, we used the Seahorse XF metabolic analyzer to measure the extracellular acidification rate (ECAR) over 2 hrs as a test for the rate of lactate production. Cells were exposed to perturbations, including the addition of glucose (to increase overall glycolysis), oligomycin (to inhibit mitochondrial ATP synthase and force maximal glycolysis), and 2-Deoxy Glucose (2D-G, a non-usable glucose analog to shut down glycolysis). We found that ISL2 KO cells had significantly reduced rates of ECAR under baseline conditions and upon each perturbation (**Figure 5D**). To further assess that this reduced ECAR is due to reduced consumption of glucose, we cultured WT and KO cells in **tritiated glucose (** 5-^3^H-Glucose), which is readily metabolized and yields tritiated water (^3^H_2_O) during the sixth step of glycolysis (**Figure 5E**, left panel). ^3^H_2_O diffuses out of cells and can be directly measured from the culture media. The KO cells exhibited significantly less ^3^H_2_O release (**Figure 5E**), further establishing that ISL2 loss results in reduced glycolytic flux.

Since *ISL2* KO cells proliferate at a faster rate, we hypothesized that the increased bioenergetic demands in these cells might be met by increased mitochondrial respiration. Accordingly, using the Seahorse XF platform, we found that ISL2-depleted cells have a significantly higher oxygen consumption rate (OCR) that WT cells, indicating increased mitochondrial respiration (**Figure 5F**). Furthermore, we observed higher MitoSox Red and MitoTracker Green staining by flow cytometry, indicating that the KO cells have higher overall mitochondrial activity and mitochondrial content, respectively (**Figure 5G**).

Given these metabolic shifts, we next sought to examine whether carbon source utilization is altered upon ISL2 depletion. In this regard, our transcriptomic analyses (**Figure 4**) indicated that ISL2 loss might increase lipid and fatty acid (FA) metabolism. Therefore, we profiled steady-state levels of ~37 major FAs in WT and ISL2 KO cells using liquid chromatography-mass spectrometry (LC-MS) analysis. The data showed that the KO cells had significantly higher levels of the essential FAs, linoleic acid, and α-linolenic acid (**Figure 5H**). ISL2 KO cells also had an elevation in polyunsaturated FA (PUFAs), including arachidonic acid and its multiple derivatives that are precursors of prostaglandins. Prostaglandin is a critical signaling lipid molecule that may also act as a ligand for PPARγ, a master regulator of FA and lipid metabolism. Notably, we observed significant upregulation of PPARγ at both the RNA (**Figure 4C**) and protein levels (**Supplementary Figure 1G**) in ISL2 KO cells. The steady-state increases in fatty acid species suggest that ISL2 KO cells may have increased capacity to take up FAs from the environment or decreased capacity to utilize them. To differentiate these two possibilities, we quantified lipid droplets (LD) in response to excess external FAs by BODIPY^493/503^ staining. LD are dynamic organelles that function as a storage depot for neutral lipids, including triglycerides and cholesterol esters^26^. Notably, although the KO cells had a lower basal LD content, they showed a more significant increase in LD levels compared to WT cells when incubated with excess FA-containing media (200 uM oleic Acid for 24 hr; **Figure 5I**). Conversely, upon starvation for 72 hrs, the ISL2 KO cells had marked depletion of LD (**Figure 5I**), indicating that ISL2 depleted cells have higher FA uptake and consumption capacity.

### ISL2 KO cells are resistant to glucose and glutamine deprivation but sensitive to OXPHOS inhibition

These results highlighted that ISL2 depletion reprograms PDAC cell metabolism and suggests that it may result in selective dependence on OXPHOS instead of glycolysis. Accordingly, we predicted that ISL2-depleted cells would be relatively resistant to glucose depletion, but sensitized to inhibitors of mitochondrial respiration and lipid metabolism. Indeed, crystal violet assays showed that the growth advantage of ISL2 KO was even more marked in low glucose media (**Figure 6A**). Since glutamine metabolism can be a significant alternative carbon source, we also tested the impact of ISL2 status of glutamine dependence by either glutamine deprivation or treatment with a glutamine transporter inhibitor (BPTES). ISL2 KO cells were significantly more resistant under both conditions (**Figure 6B and 5C**). Thus, the data show that metabolic reprogramming resulting from ISL2 loss renders cells less dependent glycolysis or glutaminolysis (**Figure 6C**).

**Figure 6:**
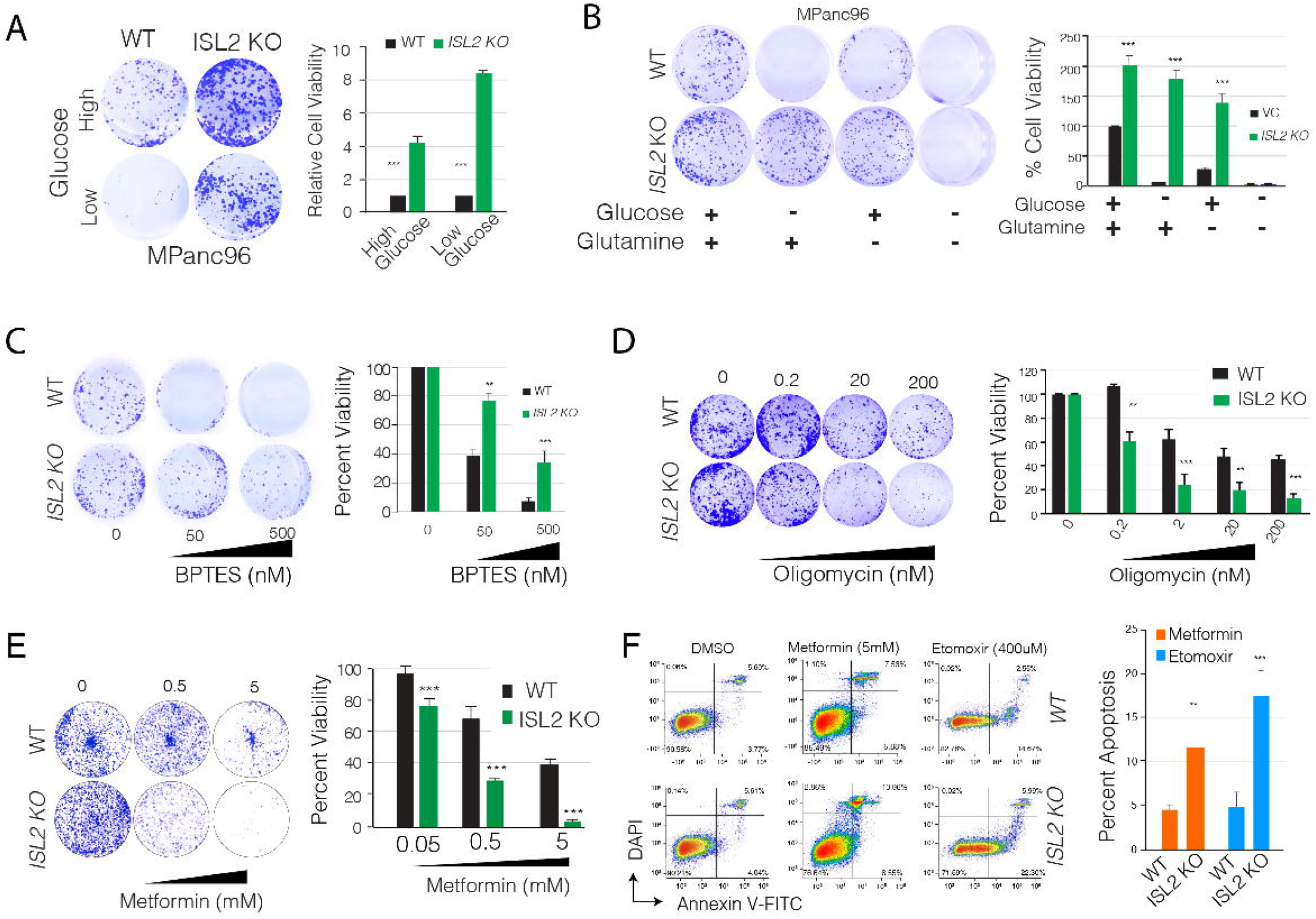
ISL2 depleted cells are sensitive to OXPHOS inhibitors. **A)** Crystal violet and Bar plots show the relative viability of WT and ISL2 KO MPanc96 cells in high (55 mM) and low glucose (5.5mM) containing media. **B)** Crystal violet and bar plots show relative viability/proliferation of WT and ISL2 KO MPanc96 cells in glucose and glutamine depleted media. **C)** Crystal violet and bar plots show the relative viability of WT and ISL2 KO MPanc96 cells at different concentrations of BPTES. **D)** Crystal violet and Bar plots show the relative viability of WT and ISL2 KO MPanc96 cells at different concentrations of oligomycin. **E)** Crystal violet and Bar plots show the relative viability of WT and ISL2 KO MPanc96 cells at different concentrations of metformin. **F)** Flow cytometry measured apoptosis rates (Annexin-V staining) in WT and ISL2 KO cells in response to metformin and Etomoxir. The bar plot shows the quantification of three independent experiments. Error bars show SEM of three or more independent experiments.

Next, we examined sensitivity to inhibitors of OXPHOS and lipid metabolism. We tested metformin, which is an FDA-approved inhibitor that modulates mitochondrial electron transport chain, and oligomycin, which inhibits mitochondrial ATP synthase. We found that ISL2 KO cells were significantly more sensitive to both of these small molecule inhibitors of mitochondrial respiration (**Figure 6D-E**). Notably, the detection of Annexin V staining by flow cytometry demonstrated that metformin causes apoptotic cell death in ISL2 KO cells (**Figure 6F**). Similar results were seen with etomoxir, which inhibits FA β-oxidation in part by blocking mitochondrial fatty acid transporter (CPT1) (**Figure 6F**). Therefore, ISL2 inactivation renders PDAC cells sensitive to inhibition of OXPHOS and FA β-oxidation. The significant apoptosis associated with a block in these processes suggests exploitable vulnerabilities in ISL2 deficient PDAC tumors.

## DISCUSSION

PDAC displays heterogeneous clinical and biological features that are not fully explained by the profile of genetic alterations in this tumor type. Here, we employed in vivo CRISPR screening to identify epigenetic regulators and transcription factors that impair the proliferation and survival of PDAC cells. This unbiased approach and subsequent validation results uncovered ISL2 as a novel suppressor of the proliferation of PDAC cells. Our functional studies indicated that ISL2 loss drives a transcriptionally-mediated metabolic switch favoring elevated OXPHOS and FAO, leading to increased dependence on these processes for tumor cell growth. Importantly, we found that epigenetic silencing of ISL2 in human PDAC specimens was associated with concordant transcriptional changes and poor outcomes. Thus, our studies reveal ISL2 status as a critical determinant of distinct transcriptional programs governing fuel source utilization and tumorigenesis of PDAC.

*ISL2* (also called *Islet 2*) encodes a TF harboring 2 LIM protein-interacting domains and a homeobox DNA binding domain (LIM/HD), which is implicated in neuronal specification^27,28^ and adipose tissue heterogeneity^29^. While ISL2 has not previously been examined in cancer, 10 of 12 related LIM/HD TFs are thought to have cancer-relevant functions^30^. In line with our studies demonstrating that ISL2 inactivation promotes tumor growth in PDAC models, genomic analysis of primary human specimens shows that *ISL2* is epigenetically silenced in >65% of PDAC tumors, and hypermethylation of *ISL2* locus correlates with low ISL2 expression and poor outcomes. Our transcriptomic analysis indicates that a significant proportion of ISL2-regulated transcriptome relates to metabolic pathways. Both the acute expression changes observed upon ISL2 inactivation in PDAC cell lines and the differentially regulated genes in ISL2-low human PDAC samples pointed to roles of ISL2 in suppressing glycolytic gene expression and enhancing OXPHOS and lipid metabolism programs. Thus, ISL2 is a tumor suppressor whose loss confers a distinct metabolic feature in PDAC.

The metabolic landscape of PDAC has been explored extensively. KRAS mutation, the defining oncogenic alteration in PDAC, is an important regulator of the metabolism of these tumors. In particular, KRAS induces high rates of glycolysis and flux into the non-oxidative pentose phosphate pathway as well as activation of glutamine metabolism, in part transcriptional upregulation of enzymes in these metabolic pathways. These pathways, in turn, support nucleotide biosynthesis and redox homeostasis, to drive PDAC growth. However, large scale metabolomic profiling studies of PDAC cell lines point to heterogeneity in the metabolic programs of PDAC subsets^31^, with distinct uses of glutamine, glucose, and fatty acids and different sensitivities to metabolic pathway inhibitors. Our results show that ISL2 loss reprograms the metabolism of PDAC cells where they upregulate mitochondrial respiration at the expense of glycolysis and become specifically hypersensitized to OXPHOS inhibition. Our findings also suggest that *ISL2* depletion is a critical molecular switch that enhances lipid metabolism in PDAC cells. Beyond *de novo* lipid synthesis, fatty acid oxidation (FAO) sustains both ATP and NADPH production, which support cell survival, growth, and proliferation^32^. In glioblastoma multiforme (GBM), acyl-CoA-binding protein (ACBP) fuels tumor growth by controlling the availability of long-chain fatty acyl-CoAs in mitochondria, promoting FAO^33^. A subset of diffuse large B cell lymphomas (DLBCL) displays enhanced mitochondrial energy transduction and greater incorporation of nutrient-derived carbons into the tricarboxylic acid (TCA) cycle^34^. Furthermore, the direct transfer of lipids from adipocytes to ovarian cancer cells induces FAO in cancer cells and promotes the migration of cancer cells to the omentum, an organ primarily composed of adipocytes^35^. While additional studies will be required to identify the critical outputs of the heightened OXPHOS and lipid metabolism in ISL2-deficient PDAC cells, our pharmacologic studies reveal specific vulnerability to the inhibition of these processes. Thus, ISL2 silencing may provide a biomarker for precision medicine approaches exploiting these metabolic shifts in PDAC.

The specific mechanisms by which ISL2 loss rewires the transcriptional and metabolic program of PDAC cells will require further investigation. Comparative transcriptome analysis in ISL2 KO cells, as well as ISL2-low primary tumors, suggests that c-MYC may be a critical factor in governing the gene expression programs in ISL2-depleted cells. While *c-MYC* were not changed upon ISL2 inactivation, the differentially regulated genes were significantly enriched for c-MYC targets, suggesting that normal levels of ISL2 may interfere with c-MYC transcriptional activity. It will also be important to identify master regulators that govern the FA and lipid metabolism in ISL2 depleted cells. To this end, PPARγ stands out as a putative candidate TF. We observed significant upregulation of PPARγ both at the RNA and protein levels in ISL2 depleted cells. PPAR family members are lipid-activated nuclear hormone receptors^36,37^, and PPARγ is a key metabolism regulator^36–38^ and has critical roles in lipid storage and FA metabolism^39^. It is notable that we also observed that the ISL2 KO cells have significantly higher levels of PUFAs, including Arachidonic acid and multiple of its derivatives that are precursors of prostaglandins (Docosa-tetra, -penta and -hexaenic acids). These FAs serve as signaling lipid molecule ligands for PPARγ ^40,41^. Thus, MYC and PPARγ appear to be good candidates for mediating the primary gene expression program in response to ISL2 depletion.

In summary, this study demonstrates that unbiased in vivo screening approaches can be exploited to identify novel transcriptional and metabolic regulators of aggressive PDAC growth^42^. Our data suggest that ISL2 is a novel tumor suppressor in PDAC and that its depletion reprograms transcriptional and metabolic state of PDAC cells. These findings provide insight into the molecular underpinnings of this lethal disease and point to OXPHOS as a potential selective vulnerability in ISL2-deficient tumors. This contributes to the growing list of cancer-specific “addiction mechanisms” due to genetic and epigenetic^43^, transcriptional^44^, and metabolic reprogramming will enable a better understanding of the disease and develop novel targeted therapies.

## Materials and Methods

### In vitro cell culture

Patient-derived tumor cells, PDX366, and MPanc96 cell line were cultured in Roswell Park Memorial Institute 1640 medium (RPMI-1640) containing 10% fetal bovine serum (FBS) and cultured in a humidified (37°C, 5% CO2) incubator.

### Generation of CRISPR sgRNA library pool and viral infection

PDX366 cell line was generated from a PDAC patient tumor as described earlier ^1^. WT Cas9 expressing lentivirus was generated in 293T cell lines by using WT Cas9 (modified from GeCKO plasmid by removing gRNA), psPAX, and pMD2.6 plasmid with 5:4:1 ratio. 10 μg total DNA was used in the presence of 30 μl of Fugene6 reagent in 10-cm plate dish that has 70 % confluency. PDX366 cell line was infected with this lentivirus for one day and then treated with 0.5μg/ml puromycin for four days. The nuclear sgRNA libraries were kind gifts from Dr. Sabatini lab (MIT) (*Wang et al.*, ^2^; Addgene Catalogue # 51047). The libraries were amplified using published protocol at Addgene. (http://www.addgene.org/static/data/08/61/acb3ad96-8db6-11e3-8f62-000c298a5150.pdf). The library pool targets 3733 transcription factors, epigenetic regulators and other nuclear function genes with ~10 sgRNA/gene. 100 non-genomic targeting control sgRNAs are included in the library. The sgRNA library expressing viruses were generated in 2× 15-cm plates by using total 20 μg DNA and the aforementioned condition. Serial dilutions of a virus were used to find the MOI of ~0.3 after selection with 5 μg/ml blasticidin for four days. Cells were harvested from 12× 15-cm plates to get at least 200× fold coverage (~8 million cells per sample) for the *in vivo* (orthotopic injection into mice pancreas) screening.

### *In vivo* CRISPR screening in an orthotopic patient-derived xenograft (PDX) model of PDAC

6-7-week-old athymic nude mice (Envigo, Indianapolis, IN) were used for *in-vivo* screening and selection. The sgRNA library, WT Cas9 expressing PDX366 cells were resuspended in 150 μL Matrigel ®Growth Factor Reduced Basement Membrane Matrix (Corning, Corning, NY). After the anesthetize, the left flank of mice was opened to exteriorize the pancreas, and 8×10^6^ PDX366 cells were injected directly into the pancreas. At this stage, one batch of cells was harvested as “Day 0” control sample. For in-vitro screening, cells were passaged every 3-4 days by 1/3 split with fresh media in 15 cm plates. At least 12 million cells were passaged each time by using 3× 15-cm plates.

Tumor volumes were monitored by MRI. MRI measurement (University of Virginia Molecular Imaging Core, Charlottesville, VA) has been performed at the conclusion of the experiment after 4 weeks. Tumors were harvested and weighed, and samples collected for further analysis. Formalin-fixed tumor samples were submitted to the University of Virginia Research Histology Core Lab for processing and H&E staining. Tumor sections were scored by a board-certified pathologist who specializes in gastrointestinal cancers. This study was carried out in strict accordance with the recommendations in the Guide for the Care and Use of Laboratory Animals of the National Institutes of Health. The animal protocol was approved by the Animal Care and Use Committee of the University of Virginia (PHS Assurance #A3245-01).

### Validation of KO efficiency by western Blot

For validation of ISL2 after initial screening, the following gRNA guiding sequences were designed, and sgRNAs were cloned to generate *ISL2* knock out cells;

sg1: GCCGGCAGAGAAGCCCGGGA,
sg2: GCGACACCCGCAGGATAAAC,
sg3: GGCAGGCCGCGTGCCACTCG.

Briefly, the oligos that have -5’CACC and -5’AAAC overhangs of the sgRNA guiding sequence were ordered from Eurofins and hybridized to get sticky end double-strand DNA for ligation. The plasmid containing the sgRNA backbone was digested with Bbs.i at 55 °C for 2 hours followed by CIP treatment at 37 °C for a half-hour. Purified vector backbone from 2 % gel (60 ng) and hybridized oligos (1 μL from 1-10 nM) were used for ligation reaction in the presence of T4 ligase.

PDX366 and MPanc96 cells were virally infected to express Cas9 and gRNA to produce stable cell lines. After 4 days of puromycin selection (2μg/ml), serial dilution was performed to generate single clones. Once the desired number of clones were obtained, lysates were prepared in RIPA buffer, and equal amounts of lysates were loaded in NuPAGE 4–12% Bis-Tris gradient gel (Invitrogen #NP0335). I-blot Nitrocellulose membrane (Invitrogen #B301002) was used for western blotting. Antibodies targeting ISL2 (Abcam #ab176576) and alpha-tubulin (Thermo Fisher Scientific # 62204) were used in the western blot.

### MTT Cell Viability

PDX366 and MPanc96 cells were seeded in a clear, flat-bottom 96-well plate (Corning) in triplicate at a density of 2-5×10^3^ cells per well. The following day, cells were treated with Oligomycin (Sigma-Aldrich #75351) and T0070907, *PPARγ* inhibitor (Selleck Chemicals LLC #S2871) for 96 hours prior to MTT (3-(4,5-Dimethylthiazolyl)-2,5-diphenyltetrazolium bromide) to determine effects of drugs on cell viability. Culture media were replaced with fresh RPMI, which has 10% FBS and 10% MTT (5mg/ml) and incubated for 4 hours in a humidified (37 °C, 5% CO2) incubator. 100 μl MTT solvent (10% SDS in 0.01M HCL) was added to each well, and cells were incubated overnight. The absorbance was read at 595 nm.

### Crystal Violet Assay

Wild-type and ISL2-KO cells were seeded in a clear, flat-bottom 12-well plate (Corning) at a density of 2×10^3^ cells per well. The following day, cells were treated with Etomoxir (Cayman Chemical #828934-41-4), Oligomycin (Sigma-Aldrich #75351), Metformin (Selleck Chemicals LLC #S1950) and BPTES (Sigma Aldrich # SML0601) for three weeks to one month to determine effects of drugs on cell viability. Culture media were replaced with fresh RPMI in the presence of inhibitors every week. Wells were washed three times with PBS, then stained for 30 minutes with crystal violet solution (0.4% crystal violet, 10% formaldehyde, 80% methanol). After staining, the solution was washed out once with PBS and water until complete removal of left-over crystal violet dye on the sides of wells. The plate was dried out overnight and imaged by the scanner.

In order to determine the response of wild-type and ISL2-KO cells to nutrient deprivation, 500 cells were plated in a 12-well plate and cultured in regular media or absence of either glucose or glutamine or both for three weeks to one month. Crystal violet staining was processed as described above.

### Annexin V Staining

In order to determine the percentage of apoptotic cells, Annexin V staining was performed and analyzed by Flow Cytometry. Cell lysates were resuspended in Annexin V binding buffer (10mM HEPES, 140mM NaCl, and 2.5mM CaCl2, pH 7.4). 5μl Annexin V conjugate (Life Technologies #A13199) was added to suspension for 15 minutes at room temperature. Stained cells were analyzed by Flow Cytometry.

### Radiotracer Metabolic Flux Studies/Substrate Competition Assay

Cells were harvested from the culture the day of assay, and 250,000 cells were seeded per well in a 24-well plate in DMEM supplemented with 10% FBS. After 3 hours of incubation, the cells were washed in PBS. Freshly made KRP assay media with 5mM glucose, 50μM acetate, 0.5mM glutamine, 1mM carnitine, and 125μM palmitate supplemented with either 3H-glucose (Perkin Elmer) for glycolysis determination or 14C-Palmitate (Perkin Elmer) for oxidative metabolism determination was applied.

For wells dedicated to the evaluation of glycolysis, wells were sealed upon addition of the assay media supplemented with 3H-glucose and incubated for 2 hours. After incubation, 1N HCl was added, and all the liquid was collected into an Eppendorf tube. The tube was placed in a scintillation vial containing an equivalent volume of water and allowed to equilibrate overnight. After equilibration, the tube was removed, scintillation cocktail fluid was mixed (Optiphase Super, Perkin Elmer) to the remaining liquid, and vials were counted on a scintillation counter (LS6500, Beckman Coulter). Counts were normalized to cell number counted from a sister plate that was seeded concurrently with the experimental plate.

For wells dedicated to the evaluation of oxidative metabolism, a CO_2_ trap of 2M NaOH in a PCR tube was placed in the wells with the assay media supplemented with 14C-palmitate. The wells were sealed and cells were incubated for 2 hours. After incubation, 2M perchloric acid was injected into each sealed well, resealed, and allowed to incubate at room temperature for 1 hour. After incubation, the NaOH was removed from the trap and placed into scintillation vials, scintillation cocktail fluid was mixed, and vials were counted on a scintillation counter. Counts were normalized to cell number counted from a sister plate that was seeded concurrently with the experimental plate.

### Mitochondrial Stress Test / Glycolysis Stress Test

Oxygen Consumption Rate (OCR) for Mitochondrial Stress Test (MST) assays and Extracellular Acidification Rate (ECAR) for Glycolysis Stress Test (GST) assays were measured using a Seahorse XF24 Extracellular Flux Analyzer. Prior to either GST or MST assays, the 24-well cell culture microplates were coated with 50 μL of poly-D-lysine (Millipore) at 10 μg/cm2/sterile water overnight. The following morning, the plate was washed once with PBS and allowed to dry.

GST media was made using serum-free DMEM without glucose, glutamine, pyruvate, sodium bicarbonate (Sigma-Aldrich) adjusted to pH 7.4. MST media was made using unbuffered, serum-free DMEM adjusted to pH 7.4 (Life Technologies). For both assays, 100,000 cells were plated in at least triplicate for each condition the day of the experiment in 100 μL of GST media per well for GST assay and 100 μL of MST media for the MST assay. Cells were then spun at 500 RPM for 1 min and supplemented with 575 μL of GST media or MST media and immediately placed into the Seahorse Analyzer to begin assay. During the GST assay, glucose (BD), oligomycin (Millipore), and 2-deoxyglucose (Chem-Impex) were injected to a final concentration of 10mM, 2 μM and 100mM, respectively over a 100-minute time course. For the MST assay, oligomycin, BAM15 (Cayman) and Rotenone and Antimycin A (Sigma) were injected to a final concentration of 2 μM, 10 μM, 1 μM and 2μM, respectively over a 100-minute time course. At the end of each experiment, each assay was normalized to cell number counted from a sister plate that was seeded concurrently with the experimental plate.

### Lipid uptake and lipid droplet analysis

Wilt-type and *ISL2* KO cells were plated in micro cover glasses (VWR cat. no. 48368-062) at a density of 2×10^4^ in regular growth conditions. After 48 hours, cells were stained by BODIPY™ FL C_12_ (4,4-Difluoro-5,7-Dimethyl-4-Bora-3a,4a-Diaza-*s*-Indacene-3-Dodecanoic Acid) (Thermo Fisher Scientific #D3922) to determine the amount of lipid droplet in regular conditions. We added 200μM Oleic Acid conjugated with BSA into cells and incubated for overnight. Then cells were stained by BODIPY to determine the amount of lipid droplet in response to oleic acid. In order to understand how they respond to starvation after oleic acid loading, we washed out media with PBS for three times to rid of oleic acid and then replace media with glucose-free media and incubated cells for 48 hours to determine the relative levels of lipid droplets. Images were captured by using Zeiss LSM 700 confocal microscopy, and the area of lipid droplet was measured by using Image J.

### Organoids xenografts

We generated 3D organoids lines from the pancreas of Pdx1^Cre^:Kras^LSLG12D^ C57Bl/6J mice. These mice were subjected to acute pancreatitis by intraperitoneal injections of cerulean, and the organoids were isolated from the PanIN precursor lesions from the pancreas. To genetically downregulate *Isl2*, we used LentiV2 constructs that express the SpCas9 and two different *Isl2* targeting guide RNAs. We generated the stable Kras^LSLG12D^ organoid lines with two different lentiviruses that target the Isl2 gene and a control guide that targets bacterial Beta-galactosidase (LacZ) gene. Subsequently, we selected the infected organoids using puromycin (2ug/ml) for a week and validated the expression of *Isl2* using qRT-PCR (Primers: Isl2_F:gcagcaacacagtgacaagg and Isl2_R ggtacgtctgcacctcgact). To determine the tumorigenicity of these organoids, we injected these three different organoid lines 4 times per each line in the subcutaneous fat of SCID mice. We injected 2 million cells from organoids per each injection and monitored for tumor growth for 6-8 weeks. We sacrificed the whole cohort once a tumor reached more than 1000mm3.

Finally, after sacrificing the whole cohort, we measured the tumor volume using Vernier calipers and tumor mass using a sensitive balance. The tumors were processed for Immunohistochemistry by fixing the tumors using 10% formalin and embedding them in paraffin. Haematoxylin and Eosin staining was performed on these tumor sections, and a blinded analysis was performed with an expert pathologist for grading these tumors.

### Targeted amplification of CRISPR/sgRNA library and sequencing

Tumors from mice and in vitro cultured cells were harvested after 4 weeks. Entire tumors and all cell pellets were used to obtain genomic DNA. Briefly, tumor samples were minced into small pieces and lysed with 8 ml SDS lysis buffer (100 mM NaCl, 50 mM Tris-Cl pH 8.1, 5 mM EDTA, and 1% wt/vol SDS). Cell pellets were processed in a similar way. Mincer tumor samples or cell pellets were treated with 100 μL proteinase K (20 mg/ml) at 55 °C for overnight incubation. The next day, entire lysis solutions were used in EtOH precipitation, and genomic DNA pellets washed with 70% EtOH twice. Pellets were resuspended in RNase-containing water and quantified by Nano-drop. For each DNA sample, 100 μg genomic DNA was used for the 1^st^ PCR reaction. We run 10 separate PCR reactions with 10 μg DNA in a single PCR tube. We used the same outer *Forward Primer* and outer *Reverse Primer* from Sabatini gRNA library-specific primers for all of the samples. These primers are different from GeCKO Array *For* and *Rev* primers. Q5-high Fidelity 2× master mix was used as polymerase from NEB (# M0429L). PCR condition for the 1^st^ PCR was; 98 °C for 30 sec, 18× (98 °C for 10 sec,63 °C for 10 sec,72 °C for 25 sec), 72 °C for 2min. After the 1^st^ PCR, all reactions were combined (10× 100 μL) in one single Eppendorf tube and vortexed well. For the 2^nd^ PCR, 5 μL PCR reaction mix from the 1^st^ PCR step was used in 100 μL total PCR reaction. PCR conditions for 2^nd^ PCR were: 98 °C for 30 sec, 24× (98 °C for 10 sec,63 °C for 10 sec,72 °C for 25 sec), 72 °C for 2min. In the 2^nd^ PCR, each sample was amplified with specific forward primers that have 6-bp barcode sequence for demultiplexing of our reads during next-gen sequencing and common reverse primer. In this setting, custom sequencing and custom indexing primers for Illumina Sequencing were used. All primer sequences used for library preparation and next-gen sequencing are listed in **Table** below. The entire solution from 2^nd^ PCR was loaded on 2% gel, and the bands around 270 bp were cut and cleaned with the Qiagen gel extraction kit (a faint band above 270 bp was noticed, likely due to carrying over of primers from 1^st^ PCR reaction). Purified PCR products were quantified by using Qubit (Invitrogen), and equimolar amounts of each PCR fragment were mixed and used for subsequent high-throughput sequencing (40 nM DNA in 20 μL). The library was sequenced on several Illumina Miseq platform to get an average 10 million reads for each sample.

**Table:**
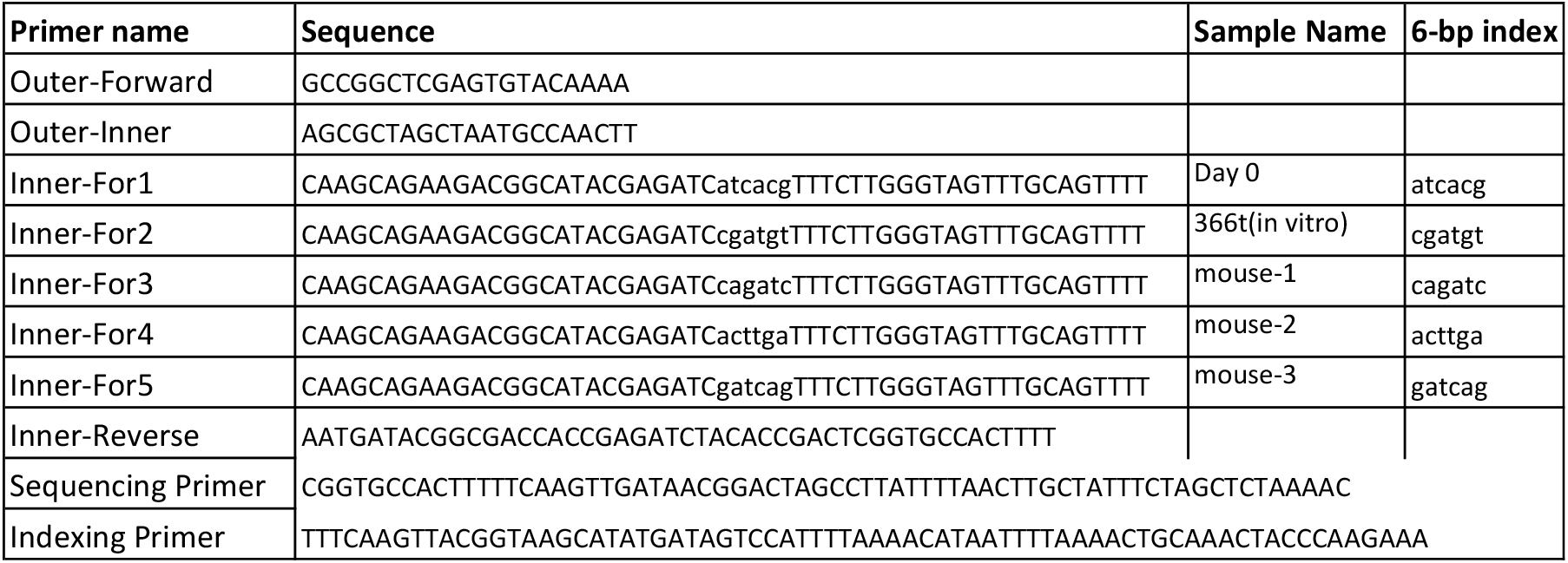
Primers used for library preparation and next-gen sequencing.

### Data analysis for CRISPR/Cas9 screening

Sequencing reads from CRISPR/Cas9 screenings were first de-multiplexed with cutadapt (v. 1.8.3). Total length 56 nt (sequencing barcode and sample barcode) were supplied to the program with the requirements that at least 36 nt of this barcode had to be present in the read so that it can be assigned to an individual tumor isolated from the PDX model. More than 99% of reads were assigned to one of the four in-vivo samples: three tumor models (further referred to as control) and cells from the day of injection (further referred to as Day 0). After de-multiplexing and removing sequencing and sample barcodes, the abundance of each sgRNA was assessed and normalized among samples with the use of MAGeCK v. 0.5.2. About 87% of the reads contained correct sgRNA sequences.

Downstream data analysis was performed in RStudio v. 0.99.484 with R v. 3.3.0 following previous publications ^2^ with slight modifications. We performed the following analysis to identify genes whose depletion positively or negatively altered the overall cellular fitness. The first step of this analysis was to calculate the relative abundance of each sgRNA by comparing normalized counts for each sgRNA between control and Day 0. Resulting numbers were log-transformed, giving log fold change (LFC) of abundance for each sgRNA. The second step of the analysis was to assign to each gene a CRISPR viability score, which is defined as a median of z-score transformed LFC of the relative abundance of sgRNAs targeting the particular gene. In the third step, we log-transformed a median of CRISPR viability score calculated for the three controls as gene enrichment score. The significance of enrichment or depletion was calculated for each gene by a Kolmogorov-Smirnov test between z-scores of sgRNAs targeting this gene and control sgRNAs.

### RNA-Seq

Total RNA was purified by using RNeasy mini kit (Qiagen #74104) by following the company's instructions. mRNA was isolated by using NEBNext Poly(A) mRNA Magnetic Isolation Module (New England Biolabs # 7490S). RNA-Seq libraries were prepared using the NEBNext Ultra Directional RNA Library Prep Kit for Illumina (New England Biolabs # E7420S) by following the company's protocol. A qubit measurement and bioanalyzer were used to determine the library quality.

### Data analysis for RNA-Seq

Overall sequencing data quality was examined using FastQC (v. 0.11.5). RNA-Seq data were aligned to the human reference genome (hg19) using HISAT2 ^3^ (v. 2.1.0) with the default paired-end mode settings. The resulting sam files were sorted by the read name and converted to bam file using samtools ^4^ (v. 1.9) sort command. Then the bam files were sorted by mapping position and indexed using corresponding samtools commands. The sorted and indexed bam files were first converted to bigwig files for visualization in the UCSC Genome Browser (https://genome.ucsc.edu/) to avoid technical alignment errors. Next, RNA-Seq data were quantified against gencode (v27lift37) annotation using Stringtie ^5^ (v. 1.3.4d) with the default settings. After obtaining the gene count matrix from Stringtie, we imported it into R and normalized the data following the pipeline of DESeq2 ^6^. Specifically, to ensure a roughly equal distribution for all the genes across samples, we used rlog transformation to stabilize expression variance for genes with different expression levels. Then samples were clustered according to Euclidean/Poisson distances to make sure replicates are clustered together. By calling the DESeq function, we determined genes with significant expression changes between the *ISL2* WT and KO samples thresholding at an adjusted P-value of 0.01. Heatmaps were produced using the pheatmap package. All other plots were generated using ggplot2. Gene set enrichment analysis ^7^ were performed using the GSEA website (http://software.broadinstitute.org/gsea/index.jsp) and the stand-alone program

### Data analysis for three publicly available PDAC studies

Methylation, RNA-Seq, whole-exome sequencing, and survival data for the TCGA-PAAD project were obtained using the TCGAbiolinks package ^8^. Methylation, mutation, and gene expression data for the PACA-CA and PACA-AU projects were downloaded from the International Cancer Genome Consortium data portal (https://icgc.org/). Since all the three studies used Illumina Methylation 450K array to profile DNA methylation and similar analysis pipelines were implemented for quantification, the 32 probes annotated for *ISL2* were selected for a direct comparison between normal tissues adjacent to the tumor area and tumor tissues. Two patients, one from TCGA-PAAD and the other from PACA-AU, had tumor and metastasis tissues available, methylation levels at ISL2 probes for which were also examined. Average methylation across all the 32 probes was calculated and compared using Student’s t-test in R.

For gene expression data generated on different platforms and analyzed using different pipelines, we employed a cross-study/platform normalization method (MC method) ^9^ to make gene expression values comparable across studies. Specifically, gene expression values were normalized within each sample and median centered within each study before merging across studies. Patients with the top and bottom quartiles of *ISL2* expression were compared for their survival. The log-rank test was used to compute the survival analysis P-value. Survival analysis further separated by patients’ gender was also performed using the same test. With RNA-Seq data available for 178 primary tumors in the TCGA-PAAD study, we separated the samples based on their *ISL2* expression levels. Then transcriptomes for samples with the top and bottom quartiles of ISL2 expression were compared using DESeq2 ^6^. An adjusted P-value cutoff of 0.05 was used to identify significantly up- and down-regulated genes in *ISL2*-low tumors. Gene set enrichment analysis was performed as previously indicated. All the plots were generated using ggplot2. The relevant statistical tests were performed using corresponding R functions.

