## Supplementary Figures for "ISL2 is an epigenetically silenced tumor suppressor and regulator of metabolism in pancreatic cancer"

### Suplementary Figures

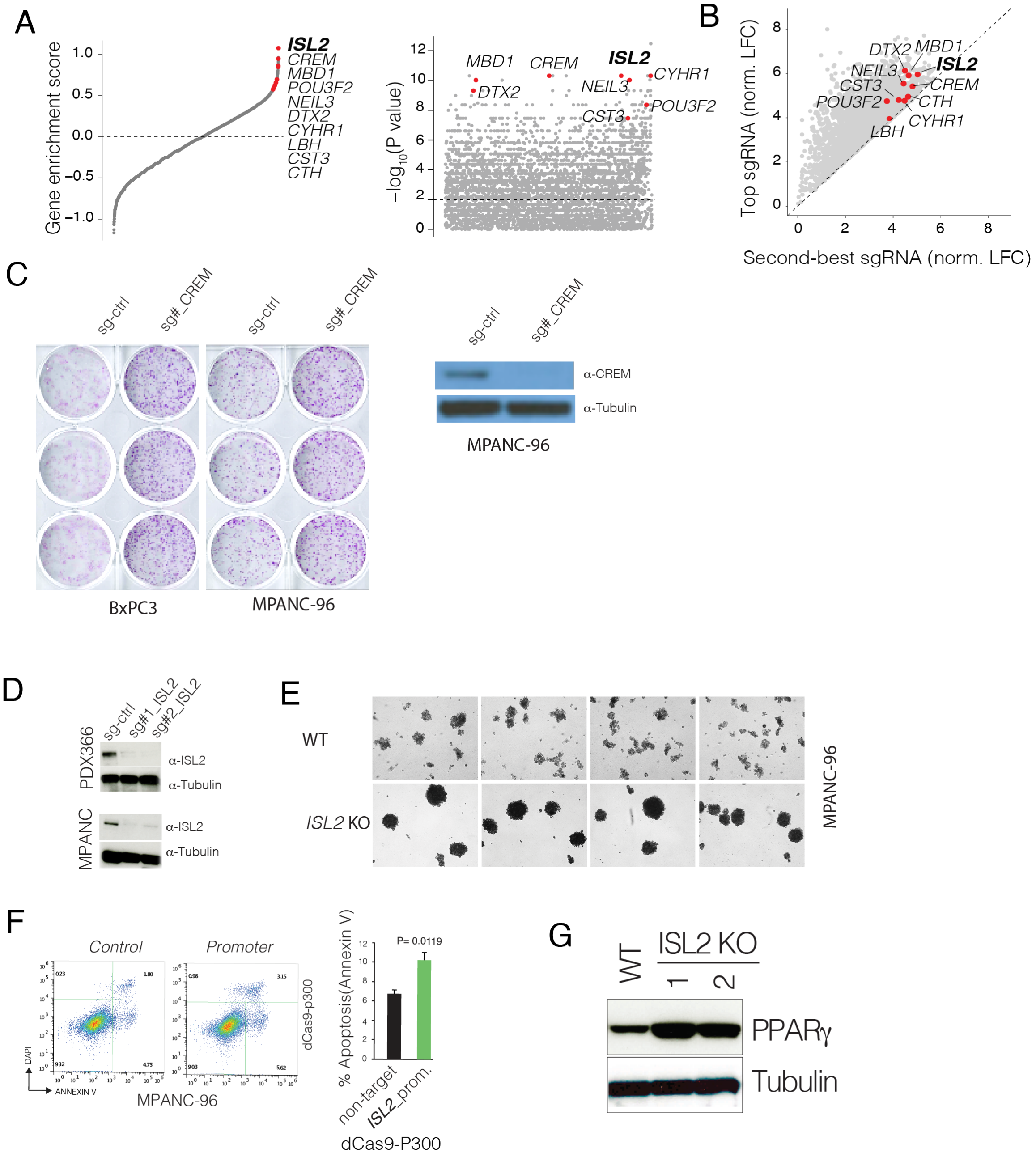


**Supplementary Figure 1: A)** Dot plot shows CRISPR viability scores (right panel) significance of gene enrichment for all genes. **B)** Dot plot shows enrichment levels of best and second best sgRNAs for indicated genes. **C)** Crystal violet assay shows relative cell proliferation levels of VC and CREM targeting BcPC3 and Mpanc96 cells. Western blot shows CREM protein levels in cells expressing control and CREM targeting sgRNA. **D)** Western shows ISL2 and tubulin levels in control and ISL2-targeting sgRNAs in PDX366 and MPanc96 cells. **E)** Images show relative levels of 3D-spheres from WT and ISL2 KO MPanc96 cells. **F)** Flow cytometry measured apoptosis (Annexin-V staining) levels in cells expressing control and ISL2-promoter targeting sgRNAs together with dCas9-P300. **G)** Western blots show PPAR-g protein levels. Tubulin levels are show as a loading control.
